# Clinical experiences with the use of oxytocin injection by healthcare providers in a South-Western State Nigeria: A Cross sectional study

**DOI:** 10.1101/474114

**Authors:** Chioma Stella Ejekam, Ifeoma Peace Okafor, Chimezie Anyakora, Ebenezer A. Ozomata, Kehinde Okunade, Sofela Ezekiel Oridota, Jude Nwokike

**Affiliations:** Department of Community Health, Lagos University Teaching Hospital, Lagos Nigeria.; College of Medicine, University of Lagos, Nigeria.; Promoting the Quality of Medicines Program, U.S. Pharmacopeial Convention.; Department of Obstetrics and Gynaecology, Lagos University Teaching Hospital, Lagos Nigeria.

**Author notes:** Corresponding Author Chioma S. Ejekam. These authors contributed equally to this work (Chioma Stella Ejekam, Ifeoma Peace Okafor, Chimezie Anyakora, Ebenezer A. Ozomata). These authors also contributed equally to this work (Kehinde Okunade, Sofela Ezekiel Oridota, Jude Nwokike).

**Keywords:** Oxytocin, clinical-experiences, oxytocin-quality, healthcare-providers, Pharmacovigilance and Lagos State

## Abstract

**Background:** Post-Partum Hemorrhage (PPH), is a leading cause of maternal mortality in Nigeria and most low and middle income countries(LMIC). The World Health Organization(WHO) strongly recommends oxytocin as effective, affordable and the safest drug of first choice in the prevention and treatment of PPH in the third stage of labor. However, there are concerns about its quality. Very high prevalence of poor-quality oxytocin, especially in Africa and Asia has been reported in literature. Excessive and inappropriate use is also common with oxytocin in low-resource settings.

**Objective:** To assess clinical experiences with quality of oxytocin used by healthcare providers in Lagos State Nigeria.

**Methods:** It was a descriptive cross-sectional study done in 2017. Seven hundred and five respondents (doctors and nurses) who use oxytocin for obstetrics and gynaecological services were recruited from 195 health facilities (public and registered private) across Lagos State. Data collection was quantitative, using a pretested self-administered questionnaire. Data analysis was done using IBM SPSS version 21. Statistical significance was set at 5%(p<0.05). Ethical approval was obtained from Lagos University Teaching Hospital Health Research Ethics Committee. Funding support was provided by the Promoting the Quality of Medicines, a program funded by the U.S. Agency for International Development and implemented by the U.S. Pharmacopeia Convention.

**Results:** Only 52 percent of the respondents knew oxytocin should be stored at 2°C to 8°C. About 80% of the respondents used oxytocin for augmentation of labor; 68% for induction of labor, 51% for stimulation of labor and 78% for management of PPH. Forty-one percent used 20IU and as much as 10% used 30IU to 60IU for management of PPH. About 13% of the respondents have experienced use of an ineffective brand of oxytocin in their practice. Just over a third (36%) of the respondents had an available means of documenting or reporting perceived ineffectiveness of drugs in their facility. Of these, only about 12% had pharmacovigilance forms available in their facilities to report the ineffectiveness.

**Conclusion:** The inappropriate and inconsistent use of oxytocin especially overdosing likely led to the spuriously high perception of medicine effectiveness among respondents. This is also coupled with lack of suspicion of medicine ineffectiveness by clinicians as a possible root cause of poor treatment response or disease progression. Poor knowledge of oxytocin storage and consequent poor storage practices could have contributed to the ineffectiveness reported by some respondents. There is need for the establishment of a unified protocol for oxytocin use with strict compliance to the guidelines. Continued training of healthcare providers in medicines safety monitoring is advocated.

## Background

Poor maternal and child health indices have remained a recurring public health challenge in Nigeria. Obstetric haemorrhage especially post-partum haemorrhage (PPH) is a leading cause of maternal mortality in Nigeria [1,2]. According to the world health organisation (WHO) post-partum haemorrhage (PPH) is defined as a blood loss of 500 ml or more within 24 hours after birth and it is said to affect approximately 2% of all women who give birth [2]. PPH is associated with nearly one quarter of all maternal deaths globally[2]. In 2015 Nigeria and India accounted for approximately 58,000 maternal deaths, which is over one third of the global maternal deaths[3]. Fortunately, deaths from PPH are preventable. WHO strongly recommends Oxytocin as an effective, affordable and the safest drug of choice in the prevention and treatment of PPH[2]. Oxytocin is also used intrapartum for induction, stimulation and augmentation of labour when medically indicated and where benefit outweighs the risk[4,5]. Oxytocin is named one of the 13 life-saving commodities by the UN Commission within the continuum of care to effectively address the avoidable causes of death during pregnancy and childbirth[6]and is included in the WHO Model List of Essential Medicines[7].

However, there are concerns about the quality of oxytocin available. Oxytocin requires stable cold chain from the point of manufacture to point of use to maintain its quality[8]. It is recommended to be stored in the refrigerator at 2° to 8° Celsius while some have extended storage up to 20° and 25°C within a few days”[8]. A major problem of oxytocin relates to heat-related degradation, inappropriate storage in the supply chain and at the health facilities. In most low-income countries these storage conditions are usually very difficult to maintain[9,10]. Surveillance studies have shown high prevalence of poor-quality oxytocin, particularly in Africa and Asia.[10–12] Most common problems were insufficient or no active ingredient[10,11]. Safe medicines supply is fundamental to public health[13]. Poor-quality medicines have the greatest potential for harming the health of consumers, with far-reaching consequences, which include: treatment failure, adverse drug reactions, economic hardship, health problems, and death[13]. Poor quality uterotonics in circulation have dire consequences. Apart from increased maternal mortality, It could also lead to performing surgical procedures that could have been prevented[14].

In a recent study in Nigeria, the quality audit of oxytocin injections in circulation showed an alarming failure rate, up 74% of sampled oxytocin injection failed quality test[10]. Despite this evidence and concerns around poor-quality medicines, epidemiologic data around quality of medicines is still spare and poor. Many healthcare providers again, do not generally suspect medicines as a cause of disease progression and a contributor to treatment outcome. Reports have it that obstetricians in Sub-Saharan Africa often give three vials of oxytocin to ensure they get equivalent of at least one dose as prevention of PPH with one vial of oxytocin is difficult.[14] This study serves as a sequel to the quality audit of oxytocin injections in Nigeria and seeks to assess the clinical experience of health care providers in Lagos State Nigeria with the quality of oxytocin injection used. It tried to assess what healthcare providers know about oxytocin injection, how they use it, what their clinical experiences were with use and their perceived effectiveness or ineffectiveness of the medicines. Effectiveness of oxytocin in this study is the ability of the oxytocin injection used, to achieve the desired contraction within the recommended dose for a specific indication.

## Materials and methods

### Study population

A descriptive cross-sectional study was carried out to assess the clinical experience of healthcare providers in Lagos State on the use of oxytocin. The study population consisted of practicing doctors and nurses working in either public or private facilities in Lagos State. To participate in the study, respondents have to be employed in registered public or private health facilities in Lagos State that offered obstetrics and gynaecological services and use oxytocin in their practice.

### Sample size determination

The sample size was determined using Cochrane’s formula considering the following criteria: standard normal deviate at 95% confidence interval; 5% accepted error of margin and proportion of reported effectiveness(52.5%) of another uterotonic from a previous study in Nigeria[15].

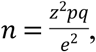

where:

*n* = minimum required sample size

*z* = standard normal deviate a 95% confidence interval = 1.96

*e* = accepted error of margin = 5%

*p* = proportion of reported effectiveness of misoprostol from a previous study in Nigeria^20^ = 52.5%

*q = 1-p*

thereby:

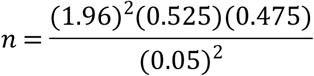

*n* = 384

Making provisions for a 30-percent nonresponse rate (to a self-administered questionnaire), the minimum calculated sample size will be 499.

## Sampling Technique and Selection of Respondents

Multistage sampling was used to select public and private healthcare facilities from each of the five administrative divisions in Lagos State.

### Stage 1: Selection of Local Government Areas (LGAs) from the five administrative divisions

A simple random sampling with ballot paper was used to select four LGAs from Ikeja administrative division; two LGAs from Badagry; two from Lagos Island; one LGA from Epe and one LGA from Ikorodu. This amounted to 10 LGAs from a total of 20 LGAs in Lagos State.

### Stage 2: Selection of public and private facilities

The three tertiary health facilities that provide obstetrics and gynecological services in Lagos State were purposively selected. Every secondary level public healthcare facility/general hospital and comprehensive primary healthcare centre (PHC) in the selected LGAs was included for recruiting respondents from the public health sector.

### Stage 3: Selection of the private health facilities

Using the list of registered private hospitals per LGA as provided by the Lagos State Ministry of Health, 15 registered private healthcare facilities that offer obstetrics and gynaecology services were selected by systematic sampling per LGA. This came to a total of 150 private health facilities from the 10 LGAs.

### Stage 4: Selection of healthcare providers

All registered doctors and nurses in the health facilities selected who met the inclusion criteria and signed the informed consent form confirming willingness to participate were included. To ensure representation, based on the proportion of doctors to nurses in the public and private sector according to the Human Resources for Health indices in Lagos State, 60 percent of respondents recruited in the study were from the private health sector, and 40 percent were from the public health sector. The doctor-to-nurse ratio of 1:2 was used to recruit respondents from both the public and private sectors.

Overall, seven hundred and five(705) respondents (doctors and nurses) who use oxytocin were recruited from 195 health facilities, which included from the public sector - 3 tertiary facilities; 10 General hospitals; 32 Comprehensive Primary Health Care facilities and private sector - 150 private health facilities across the 5 administrative divisions of Lagos State. This is shown in Table 1.

**Table 1:**
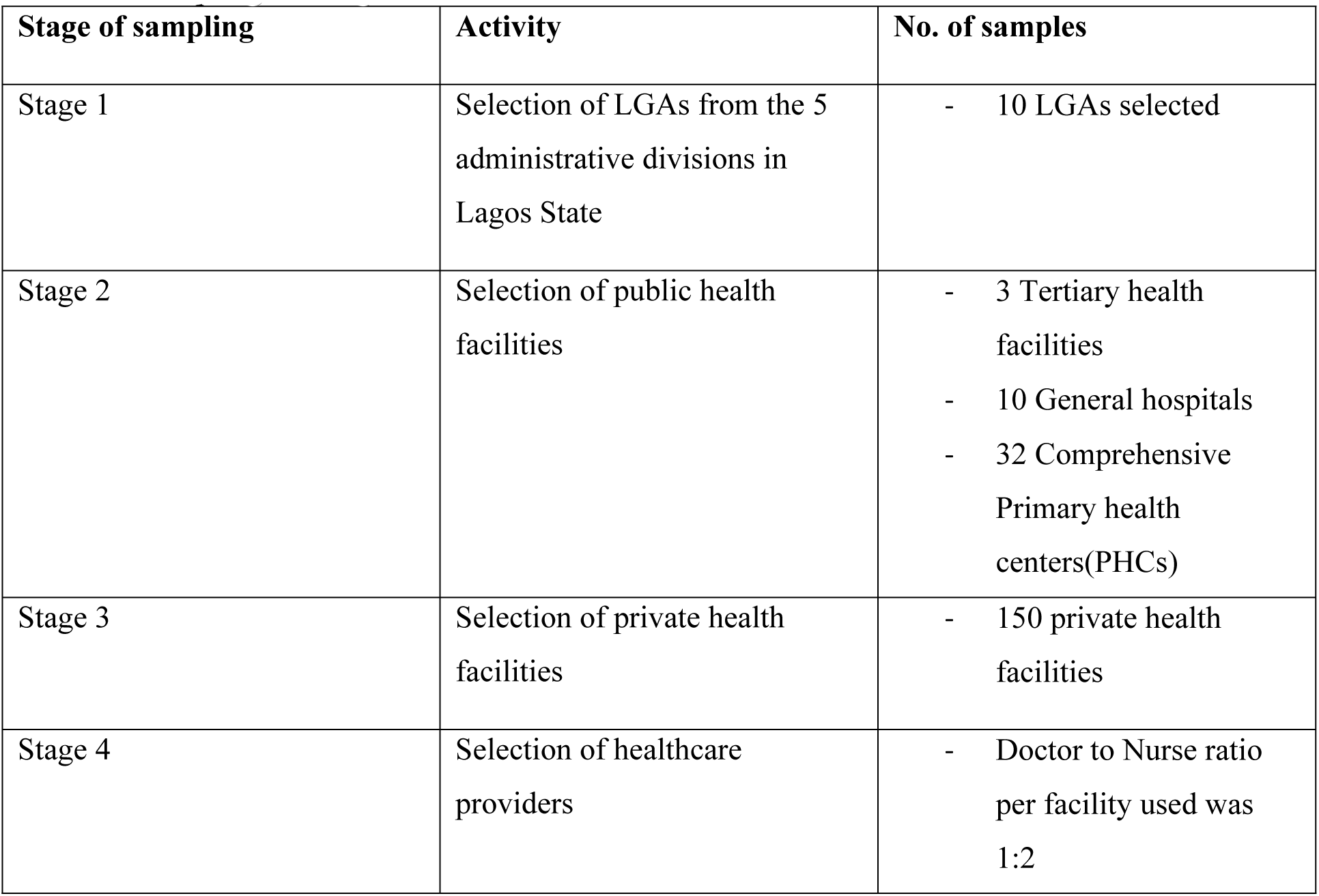
Sampling technique.

| Stage of sampling | Activity | No. of samples |
| --- | --- | --- |
| Stage 1 | Selection of LGAs from the 5 administrative divisions in Lagos State | - 10 LGAs selected |
| Stage 2 | Selection of public health facilities | - 3 Tertiary health facilities<br>- 10 General hospitals<br>- 32 Comprehensive Primary health centers(PHCs) |
| Stage 3 | Selection of private health facilities | - 150 private health facilities |
| Stage 4 | Selection of healthcare providers | - Doctor to Nurse ratio per facility used was 1:2 |

Table 1 gives a description of every stage of sampling technique

### Data collection technique and management

Quantitative data was collected using a pretested self-administered questionnaire which was developed following a literature review and incorporated expert contributions, reviews, and opinions. The questionnaire was pretested among 20 clinicians (doctors and nurses) who met the inclusion criteria and were from facilities in the LGAs not selected for this study. The questionnaire sought information on sociodemographic characteristics of respondents and occupational history of respondents; general obstetric knowledge; clinical experience with Oxytocin use. The main outcome measures were proportion of healthcare providers with good knowledge of oxytocin storage, pattern of oxytocin usage and dosing by healthcare providers, the proportion of oxytocin perceived to be effective and ineffective by healthcare providers and proportion of healthcare providers who document or take action concerning perceived ineffectiveness of oxytocin in their clinical practice. Data entry, cleaning and analysis were done using IBM Statistical Package for Social Sciences (SPSS) version 21. Data was presented in frequency tables, simple proportions and inferential statistics was done with chi-square and statistical significance set at p<0.05. The mean and standard deviation were used to summarize quantitative variables that were normally distributed, while median and interquartile ranges (IQRs) were used for those that were found not to be normally distributed. Since the responses in the study were self-reported, there was possibility of recall bias and social desirability bias. However, setting the respondents recall period to the past one(1) year helped minimize recall bias. Many options were offered on each question for respondents to choose from and also the questionnaire is self-administered without respondent’s personal details(anonymity), these will reduce chances of social desirability bias. In addition, every respondent who participated in this study only had a few minutes to fill in the questionnaires. The questionnaires were not sent in advance hence no time to prepare for the ‘apparently correct’ answer, the questionnaire clearers stated that there were no right or wrong answers, they study only wanted to assess the current practice. Again because it was a self-administered questionnaire, there was also the possibility of none response bias, which was minimized here by almost doubling the minimum calculated sample size. Ethical approval was obtained from the Health Research and Ethics Committee – Lagos University Teaching Hospital, Lagos Nigeria (HREC assigned no. ADM/DCST/HREC/APP/1800). Formal consent was obtained from each respondent.

## Results

Seven hundred and five(705) respondents were participated in the study. Table 2 gives the socio-demographic characteristics of the respondents. They were mostly in within the 30 to 40 years age bracket (41.4%) with a mean age of 36.3±10.4. There were more females (71.6%) and nurses (61.0%). More respondents came from the private sector (62.1%). Majority of the respondent had ten years or less working experience (64.5%) with a median of 7.5years. Nearly all of the respondents (92.9%) had received some form of training on oxytocin.

**Table 2:** Socio-demographic characteristics of respondents (n=705)

| Characteristic | Category | Frequency | Percent |
| --- | --- | --- | --- |
| Age group (years)* | <20 | 2 | 0.3 |
|  | 20–29 | 195 | 27.7 |
|  | 30–39 | 292 | 41.4 |
|  | 40–49 | 124 | 17.6 |
|  | 50–59 | 66 | 9.4 |
|  | ≥60 | 26 | 3.6 |
| Sex | Males | 200 | 28.4 |
|  | Females | 505 | 71.6 |
| Years of practice<sup>†</sup> | 1–10 | 455 | 64.5 |
|  | 11–20 | 146 | 20.7 |
|  | 21–30 | 73 | 10.4 |
|  | ≥31 | 31 | 4.4 |
| Cadre of health worker | Doctor | 275 | 39.0 |
|  | Nurse | 430 | 61.0 |
| Sector of practice | Public | 267 | 37.8 |
|  | Private | 438 | 62.1 |
| Training on oxytocin use | Yes | 655 | 92.9 |
|  | No | 50 | 7.1 |

\* Mean 36.3, SD ± 10.4
† Median 7.5, IQR (4, 15)

Table 3 gives the general obstetrics knowledge and practice of the respondent. Most of the respondent knew the correct definition of PPH (86.2%). Just over half (52.2%) of the respondents knew the proper storage place for oxytocin i.e. in the refrigerator at 2-8°C while as much as 42% stored their oxytocin on the shelves. Analysis of the indications for use of oxytocin among respondents showed that 80% of the respondents used oxytocin for Augmentation of labour, 78.2 % for Management of PPH, 68% for Induction of labour while 50.6% used it for stimulation of labour as shown in figure 1. Figure 2 shows the respondents understanding of the proper storage conditions.

**Table 3:**
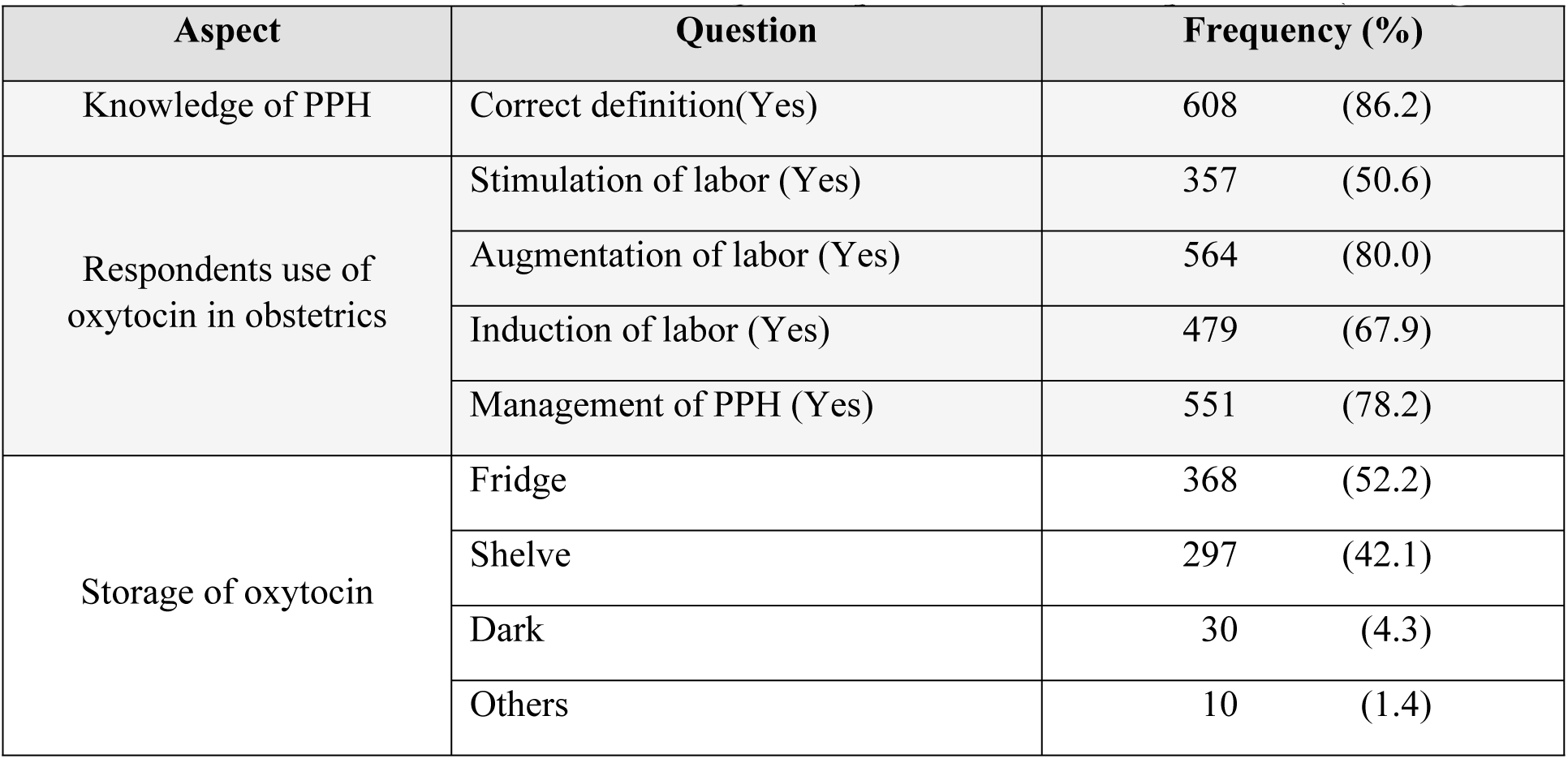
General obstetrics knowledge and practice of the respondents (n=705)

Table 3 assessed respondents’ knowledge of definition of post-partum haemorrhage, their knowledge of the recommended storage for oxytocin and what they use for in their practice.

More doctors (59.3%) than nurses (47.7%) correctly knew that oxytocin should be stored in the refrigerator and as well more respondents in government facilities (68.4%) than in private facilities (40.2%) knew the proper storage of oxytocin as shown in table 4.

**Table 4:** Assessment of knowledge of oxytocin storage by Cadre and Sector of practice of respondents.

| Question | Frequency, N=705(%) |  |
| --- | --- | --- |
| Storage of oxytocin | Doctors, n=275 (%) | Nurses, n=430 (%) |
| Refrigerator | 163 (59.3) | 205 (47.7) |
| Shelves | 93 (33.8) | 204 (47.4) |
| Dark | 14 (5.1) | 16 (3.7) |
| Others | 5 (1.8) | 5 (1.2) |
| Storage of oxytocin | Public, n=267 (%) | Private, n=438 (%) |
| Refrigerator | 182 (68.4) | 176 (40.2) |
| Shelves | 66 (24.8) | 217 (49.5) |
| Dark place | 3 (1.1) | 27 (6.2) |
| Others | 16 (6.0) | 18 (4.1) |

Table 4 further presents the analysis of the assessment of respondents’ knowledge of recommended storage place for oxytocin by healthcare provider cadre and by sector of practice.

Twenty-three percent of the respondents monitored the effectiveness of oxytocin using the frequency and duration of uterine contraction while 65% uses frequency/duration of uterine contractions and cervical dilatation as shown in table 5.

**Table 5:** Respondents’ general practice with oxytocin (n=705)

| Questions | Response | Frequency (%) |
| --- | --- | --- |
| Cadre administering oxytocin | Doctors | 165 (23.4) |
|  | Nurses | 46 (6.6) |
|  | Both | 494 (70.0) |
| Responsibility for procurement of oxytocin | Clients | 95 (13.5) |
|  | Facility | 610 (86.5) |
| Indicator for monitoring effectiveness of oxytocin in labor | Correct frequency and duration of uterine contractions | 165 (23.4) |
|  | Cervical dilatation | 73 (10.4) |
|  | Both of the above | 461 (65.4) |
|  | None of the above | 3 (0.4) |
|  | Others | 3 (0.4) |

Table 5 shows the cadre administering oxytocin in respondents’ clinical practices, who procures and their indicator for monitoring effectiveness of oxytocin used during labor.

Table 6 summarizes the respondent’s indication for use and dosage of oxytocin for various obstetric indications. About 48% of the respondents indicated they use 10IU of oxytocin for stimulation/augmentation of labour in primigravida, 24% use 5IU, 17% use 20IU, while 2.4% use other doses ranging from 30IU to 60IU. About forty-percent of the respondents use 5IU for stimulation/augmentation of labour in multiparas, 41% use 10IU, 10% use 20IU and as much as 4.4% use other doses ranging from 30IU and 60IU. Concerning the dosing of oxytocin for prevention of PPH, 41.4% of the respondents use 20IU, 33% use 10IU, 11.3% use 5IU, 4.8% use 15IU while as much 10% use doses ranging between 30IU to 60IU.

**Table 6:** Use and dosage of oxytocin for various obstetric indications.

| Dose Amount | Max dose for stimulation/<br>augmentation of labor<br>in primigravida<br>Frequency (%) | Max dose for stimulation/<br>augmentation of labor<br>in multipara<br>Frequency (%) | Dose of oxytocin for<br>prevention of PPH<br>Frequency (%) |
| --- | --- | --- | --- |
| 5IU | 169 (24.0) | 280 (39.8) | 80 (11.3) |
| 10IU | 339 (48.1) | 292 (41.4) | 230 (32.6) |
| 15IU | 57 (8.1) | 32 (4.5) | 34 (4.8) |
| 20IU | 123 (17.4) | 70 (9.9) | 291 (41.4) |
| Others (30IU to 60IU) | 17 (2.4) | 31 (4.4) | 70 (9.9) |

Table 7 shows that the majority of respondents in both public (41.4%) and private (52.3%) health facilities use a maximum dose of 10IU of oxytocin for stimulation/augmentation of labor in a primigravida. Similar responses were also noted for the same indication in the multipara in both public (41.4%) and private (41.1%) health facilities. The majority use a maximum dose of 20IU units of oxytocin for the prevention of PPH in both public (43.2%) and private (40.0%) health facilities.

**Table 7:** Use and dosage of oxytocin for various obstetric indications according to respondent’s Sector of practice.

|  | Dose Amount | Max dose for stimulation/<br>augmentation of labor in<br>primigravida<br>Frequency (%) | Max dose for stimulation/<br>augmentation of labor in<br>multipara<br>Frequency (%) | Dose of oxytocin for prevention of<br>PPH<br>Frequency (%) |
| --- | --- | --- | --- | --- |
| **Public Sector**<br>**n=267** | 5IU | 78 (29.3) | 106 (39.8) | 30 (11.3) |
|  | 10IU | 110 (41.4) | 110 (41.4) | 82 (30.8) |
|  | 15IU | 28 (10.5) | 10 (3.8) | 14 (5.3) |
|  | 20IU | 47 (17.3) | 32 (12.0) | 115 (43.2) |
|  | Others | 4 (1.5) | 7 (2.6) | 26 (9.4) |
| **Private Sector**<br>**n=438** | 5IU | 90 (20.5) | 172 (39.3) | 49 (11.2) |
|  | 10IU | 229 (52.3) | 180 (41.1) | 146 (33.3) |
|  | 15IU | 32 (7.3) | 21 (4.8) | 18 (4.1) |
|  | 20IU | 73 (16.7) | 38 (8.7) | 175 (40.0) |
|  | Others | 14 (3.2) | 27 (6.1) | 50 (11.4) |

Table 7 shows further analysis of different doses of oxytocin used by respondents per sector of practice.

Twelve popular brands of oxytocin were assessed in this survey. These brands were previously studied audited for quality [2]. The respondents’ perception of the effectiveness and ineffectiveness of these brands vary significantly. These brands were de-identified for the purpose of this research. Table 8 gives the summary of the perception of effectiveness/ineffectiveness of these brands. Overall, 98.3% have had experiences of effectiveness with the oxytocin brands while 12.6% perceive the oxytocin brands they use were ineffective as seen in Table 9.

**Table 8:** Experience of quality of oxytocin brands used in obstetrics practice (n=705)

| Brands<br>de-identified | Perceived quality of the different oxytocin brands |  |  |
| --- | --- | --- | --- |
|  | Effective<br>Frequency (%) | Ineffective<br>Frequency (%) | Don't know<br>Frequency (%) |
| A | 450 (63.8) | 29 (4.1) | 226 (32.1) |
| B | 430 (61.0) | 17 (2.4) | 258 (36.6) |
| C | 122 (17.3) | 24 (3.4) | 559 (79.3) |
| D | 602 (85.4) | 38 (5.4) | 65 (9.2) |
| E | 149 (21.2) | 33 (4.6) | 523 (74.2) |
| F | 65 (9.2) | 22 (3.1) | 618 (87.7) |
| G | 38 (5.5) | 26 (3.6) | 641 (90.9) |
| H | 48 (6.8) | 17 (2.4) | 640 (90.8) |
| I | 50 (7.1) | 19 (2.7) | 636 (90.2) |
| J | 31 (4.4) | 18 (2.5) | 656 (93.1) |
| K | 23 (3.3) | 16 (2.2) | 666 (94.5) |
| L | 38 (5.4) | 0 (0.0) | 667 (94.6) |

**Table 9:**
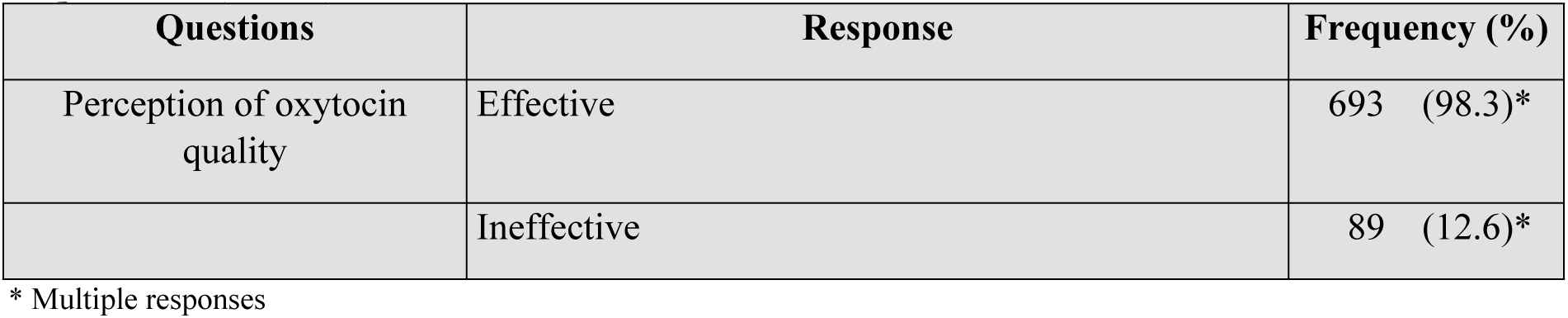
Overall experience of effectiveness and ineffectiveness of oxytocin brands used by respondents (n=705)

| Questions | Response | Frequency (%) |
| --- | --- | --- |
| Perception of oxytocin quality | Effective | 693 (98.3)* |
|  | Ineffective | 89 (12.6)* |

\* Multiple responses

Table 8 shows the respondents’ experience of effectiveness or ineffectiveness with use of each brand of oxytocin listed however, the brands were de-identified here.

Table 9 shows the pooled estimate of the effectiveness and ineffectiveness of the oxytocin brands used by the respondents.

Majority (64.3%) of the respondents have no available means in place within their facility to document and/or report experience of ineffectiveness. Of the few who have, most (61%) document it in the patient’s case note, 27% in the clinical summary and 12% in the pharmacovigilance form. In the event of oxytocin failure, 57 percent will resort to caesarean section, while 45.2 percent will change to another medicine, mainly misoprostol (40.1%). These results are summarized in table 10.

**Table 10:** Practice following oxytocin use in obstetrics (n=705)

| Question | Response | Frequency (%) |
| --- | --- | --- |
| Availability of means of documenting/<br>reporting perceived oxytocin<br>ineffectiveness | Available | 252 (35.7) |
|  | Not available | 453 (64.3) |
| Where respondents report/document<br>perceived poor quality of medicines<br>(n=453) | Case note | 154 (61.1) |
|  | Clinical summary | 68 (26.9) |
|  | Pharmacovigilance form | 30 (11.9) |
| Actions taken by respondents when the<br>maximum recommended dose of oxytocin<br>fails | Doubling the dose | 37 (5.2)* |
|  | Change the medicine | 319 (45.2)* |
|  | Caesarean section | 402 (57.0)* |

\* Multiple responses

Table 11 shows that respondents in the public sector and doctors had significantly better knowledge of oxytocin storage (p<0.001).

**Table 11:** Association between sector, cadre, and knowledge of proper storage of oxytocin(Refrigerator) (n=705)

| Question | Response | Proper storage of oxytocin |  |  |  |  |
| --- | --- | --- | --- | --- | --- | --- |
| | | Yes<br>Frequency (%) | No<br>Frequency (%) | Total | $\chi^2$ | P |
| Sector of practice | Government | 183 (68.5) | 84 (31.5) | 267 | 757.88 | <0.001 |
|  | Private | 176 (40.2) | 262 (59.8) | 438 |  |  |
| Cadre of health worker | Doctor | 159 (57.8) | 116 (42.2) | 275 | 713.34 | <0.001 |
|  | Nurse | 200 (46.5) | 230 (53.5) | 430 |  |  |

## Discussion

The healthcare system in most low income countries are weak. This situation is further exacerbated when poor quality medicines are also in circulation. Our findings suggest poor knowledge of oxytocin storage among the respondents. There was also inappropriate and inconsistent use of oxytocin with the experience of ineffectiveness of oxytocin brands used among respondents. Oxytocin is a peptitde with a highly unstable structure. The biggest obstacle to oxytocin quality is the storage and handling before patient use. The storage condition of oxytocin has been widely reported as inappropriate[16]. Oxytocin is a heat– sensitive medicine and should be kept between 2–8°C. Previous study in India documented that most physicians and nurses did not know how oxytocin should be stored[17]. An assessment in Nepal, found that only 8.6% of health facilities stored oxytocin in the refrigerator[18]. Similarly, our study, showed that only 52% of the respondents knew that oxytocin should be stored in the refrigerator while in practice this may just be much lower. A further assessment of the association between good knowledge of proper storage of oxytocin with sector of respondents’ practice revealed that about 68% of healthcare providers in the public sector and 40% in the private sector knew that oxytocin should be stored in the refrigerator. This was statistically significant with p<0.001. There was also a statistically significant difference (p<0.001) between cadre of staff and knowledge. As much as 41% of doctors and 52% of nurses did not know that oxytocin should be stored in the refrigerator.

The therapeutic dose of oxytocin for induction, stimulation and augmentation of labor for medically recommended reason is 5IU including prevention of PPH[4]. However, WHO recommends 10IU (IV/IM) for prevention and treatment of PPH[2]. Most evidence-based guidelines(U.K and Canada) in literature recommend a low dose oxytocin for induction and augmentation[20]. Our findings revealed that different doses of oxytocin (low and high) were used by healthcare providers in this study even within same facility. A very high proportion of the respondents in our study used doses beyond the maximum recommended for intra-partum use in primigravida and also in multiparas who obviously need lower doses. About 41% of the respondents used double the WHO recommended dose. Nearly 10% used doses ranging between 30IU to 60IU of oxytocin. These translate to use of two to six vials for a 10IU vial to achieve desired result of uterine contraction. This may just be an indication of failed quality, supporting the report in literature that healthcare providers in Africa often used up to 3 vials to get the desired effect of one[14]. The findings in our study are similar to reports from a previous study done in Karnataka, India[17]. This encourages wastage and diversion of limited resources for saving lives of other women and improving maternal health. It also increases the client’s healthcare spending.

In assessing respondents’ perceived effectiveness/ineffectiveness of oxytocin used in their practice, overall, up to 13% of the respondents have experienced use of an ineffective brand of oxytocin at one time or the other. Lack of suspicion of medicines quality by healthcare providers as a possible cause of disease progression or contributor to treatment outcome may have influenced this level of perceived ineffectiveness. Medicinal products are supposed to protect patients and save lives hence should be 100% effective. The findings correlate with the reports of high prevalence of poor-quality oxytocin samples in LMIC countries from laboratory assays^[11,19]^. No previous study within our search of published literature had assessed healthcare provider perceived effectiveness or ineffectiveness of oxytocin used in their clinical practice hence, posing a challenge in making comparisons.

The high level of knowledge of the correct definition of PPH is not surprising as our respondents were supposedly highly skilled healthcare providers. Similarly in Ethipoia, 82.4% of the skilled healthcare providers defined PPH correctly[21].

The pattern of indications for oxytocin use is similar to the Nepal study where majority (78%) of the health service providers used oxytocin for prevention and management of PPH while 59% used oxytocin for augmentation and induction of labour[18]. Our study is consistent with previous studies that oxytocin may be very commonly and inappropriately used for induction and augmentation of labour[20,22], almost routinely used and given contrary to guideline during labor in spite of good progress[23,24].

It was noted that majority (57%) of the respondents performed a caesarean section when the maximum recommended dose of oxytocin failed, while 5.2% doubled the dose of oxytocin used. Possible consequence of poor oxytocin quality as reported in previous studies could result to excessive and inappropriate use of oxytocin and performing unnecessary surgical procedures which could lead to avoidable complications and even death[13,14,20]. Despite these experiences only about 36% of the respondents had a system in place for documenting or reporting perceived ineffectiveness of drugs used. This again further support reports that healthcare providers often do not suspect drug quality in the course of practice and usually do not document[13].

It is possible that the inappropriate and inconsistent use of oxytocin—especially overdosing—likely led to the spuriously high perception of medicine effectiveness among respondents. This is also coupled with lack of suspicion of medicine ineffectiveness by clinicians as a possible root cause of poor treatment response or disease progression. Poor knowledge of oxytocin storage and consequent poor storage practices could have contributed to the ineffectiveness reported by some respondents.

## Strength and Limitation

There is dearth of published data on the perceived quality of oxytocin used by healthcare providers and so this study contributed to the much needed data on this topical issue especially in low and middle income countries with high maternal mortality mainly due to haemorrhage. Another strength, is the representativeness of the respondents from public and private sector and the involvement of all levels of the health system (tertiary/secondary/primary) across Lagos State. However, the study did not include middle and lower level healthcare providers such as community health officers, community health extension workers or Traditional birth attendants who also use oxytocin in their practice though not approved to use it at that level. There could be the issue of possible recall bias since the responses were self-reported. A qualitative aspect to compliment the quantitative data collected will be considered in further studies.

## Conclusion and recommendation

This study brings to the consciousness of healthcare providers in Nigeria the possible contribution of poor medicines quality to the poor maternal health risks and indices in Nigeria. It further highlighted the level of pharmacovigilance in the healthcare system and by extrapolation in various similar setting in the low and middle-income countries. Other findings include the not so encouraging level of knowledge for proper storage conditions and consequent storage practice of oxytocin and by extension the poor clinical outcomes of poor quality oxytocin. These findings align with a previous quality audit of oxytocin injections in Nigeria.

These have dire consequences. Over half of the respondent will resort to surgical procedures when the administered oxytocin is ineffective. This calls for an urgent plan to put in place a standard protocol to guide the practices in the storage and use of oxytocin. Proper reporting channels on suspected poor quality of drugs should be improved including continued education of health workers on the use of pharmacovigilance forms and ensuring availability. There is need for continuous and expanded pre-service and in-service training of healthcare providers to develop skills in drug safety monitoring including the suspicion of drug quality in the chain of events that could possibly result in poor health outcomes.

## Acknowledgements

The authors are grateful to all the respondents who participated in this study. The authors are also grateful to some staff of USP that supported this study: Olutoun Sanusi, Mopa Esuga and Adebola Adekoya.

---

**Figure 1: Indications for use of oxytocin among respondents**

